# Morphology-driven downscaling of Streptomyces lividans to micro-cultivation

**DOI:** 10.1101/159509

**Authors:** Dino van Dissel, Gilles P. van Wezel

## Abstract

Actinobacteria are prolific producers of secondary metabolites and industrially relevant enzymes. Growth of these mycelial microorganisms in small culture volumes is challenging due to their complex morphology. Since morphology and production are typically linked, scaling down culture volumes requires better control over morphogenesis. In larger scale platforms, ranging from shake flasks to bioreactors, the hydrodynamics play an important role in shaping the morphology and determining product formation. Here, we report on the effects of agitation on the mycelial morphology of *Streptomyces lividans* grown in microtitre plates (MTP). Our work shows that at the proper agitation rates cultures can be scaled down to volumes as small as 100 μl while maintaining the same morphology as seen in larger scale platforms. Using image analysis we compared the morphologies of the cultures; when agitated at 1400 rpm the mycelial morphology in microcultures approached that obtained in shake flasks, while product formation was also maintained. Our study shows that the morphology of actinobacteria in microcultures can be controlled in a similar manner as in larger scale cultures by carefully controlling the mixing rate. This could facilitate high-throughput screening and upscaling.

## INTRODUCTION

Actinobacteria produce a plethora of bioactive natural products, such as antibiotics, anticancer agents, immunosuppressants and antifungals (Barka 2016; Bérdy 2005; Hopwood 2007). In addition, these bacteria produce many industrially relevant enzymes, such as cellulases, amylases and proteases (Vrancken and Anne 2009). Streptomycetes exhibit a complex multicellular life cycle (Claessen 2014). This starts with a single spore that germinates to form vegetative hyphae, which then grow out following a process of hyphal growth and branching to produce a branched vegetative mycelium (Chater and Losick 1997). Nutrient depletion and other environmental stresses induce development, whereby aerial hyphae are formed that differentiate into chains of spores following a complex cell division event whereby ladders of septa are produced within a short time span (Jakimowicz and van Wezel 2012; McCormick 2009). In a submerged environment streptomycetes grow as mycelial networks, typically forming large pellets or clumps. From the industrial perspective, growth as pellets is unattractive, in particular because of mass-transfer problems, slow growth and culture heterogeneity (van Dissel 2014; van Wezel 2009).

High throughput (HT) cultivation methods at a small scale are highly desirable, among others to exploit the potential of newly isolated actinobacteria (Kolter and van Wezel 2016). Down-scaling of culture volumes, while maintaining key factors that influence the productivity seen in shake flasks or small scale bioreactors, is necessary to make large screening efforts rapid and economically feasible (Long 2014). However, growing streptomycetes in small cultures is challenging. Streptomycetes typically display a wide range of morphologies in submerged cultures, including dense pellets as well as large mycelial mats (reviewed in (van Dissel 2014)). After inoculation, spores germinate and produce at least two different extracellular polysaccharides (EPS), a cellulose-like polymer CslA/GlxA (Liman 2013; Petrus 2016) and a second EPS synthesized by MatAB (van Dissel 2015). Both polymers induce spore aggregation, and play a key role in pellet formation ((Zacchetti 2016) and our unpublished data). Spore aggregation promotes the formation of pellets, spatially heterogeneous structures with a largely physiologically inactive core, while the peripheral hyphae grow exponentially by tip extension and branching (Celler 2012). The relationship between pellet morphology, hydrodynamics (and oxygen supply) and production has been well studied for bioreactors (Tamura 1997; Roubos 2001; Ohta 1995) and for shake flasks (Mehmood 2012; Dobson 2008).

To successfully down-scale liquid-grown cultures, the morphology *Streptomyces* mycelia adapt in larger scale platforms (*i.e.* shake flasks or bioreactors) should be mimicked as closely as possible. The exact morphology, determined by size, density and shape, also depends on the characteristics of the environment (Wucherpfennig 2010). The hydrodynamics, in other words the characteristics of the agitated medium, is of particular importance as it influences among others the rate of fragmentation (Olmos 2013). Low agitation causes poor distribution of nutrients and reduced oxygen transfer rates, stunting growth and production, while strong agitation can cause cell death (Roubos 2001). Examples of HT cultivation platform for filamentous microorganisms have been described (Minas 2000; Siebenberg 2010; Sohoni 2012). These authors made use of shaken deep-well plates, which results in higher oxygen transfer rates than in small-volume MTPs (Duetz 2000). The Biolector system allows growth in 48 parallel 1-mL cultures (Rohe 2012; Huber 2009), which recently was successfully adapted for growth of streptomycetes (Koepff 2017).

In this work we sought to further scale down *Streptomyces* cultures to 100 μl scale. As hosts we used *Streptomyces coelicolor*, a model streptomycete for the study of development and antibiotic production (Barka 2016), and the related *Streptomyces lividans*, the preferred enzyme production host (Anné 2012). Cultures were scaled down from shake flasks to 100 μl cultures, using a digital vortex to obtain the extensive mixing required to control pellet morphology. Using whole slide image analysis, the mycelia were quantified and compared in terms of size and shape. This allowed further optimization of growth in microcultures.

## MATERIALS AND METHODS

### Bacterial strains, plasmids

*Streptomyces lividans* 66 (Cruz-Morales 2013) was used for morphological analysis and enzyme production and *Streptomyces coelicolor* A3(2) M145 was used for antibiotic production. Plasmid pIJ703, which carries the *melC1* and *melC2* genes for heterologous tyrosinase production (Katz 1983), was transformed to its host by protoplast transformation (Kieser 2000). Spores were harvested from soy flour mannitol agar plates and stored in 20% glycerol at −20°C as described (Kieser 2000). The spore titre was determined by plating serial dilutions on SFM agar plates and counting CFUs.

### Cultivation conditions

For cultivation in shake flasks, *S. lividans* was grown in 30 mL tryptic soy broth (Difco) with 10% sucrose (TSBS) in a 100 mL Erlenmeyer flasks equipped with a stainless steel spring. The flask was inoculated with 10^6^ CFUs/ ml and cultivated at 30°C in an orbital shaker with 1 inch orbit (New Brunswick) at 200 RPM. For the production of tyrosinase 25 μM CuCl_2_ was added to the TSBS medium. For antibiotic production *S. coelicolor* was cultivated in Yeast Extract - Malt Extract (YEME; (Kieser 2000)) but without sucrose (YEME0).

100 μL media with 10^6^ cfu/mL spores was added to wells of a V-bottom 96 well MTP (Greiner Bio-One, Germany). To minimize evaporation, the plate was covered with a custom moulded silicone sheet made from MoldMax40 (Materion, USA), using the 96 well plate as a mold. An AeraSeal film (Excel Scientific, USA) was added to the top for sterility, while allowing gas exchange. The combined silicone sheet and AeraSeal film were fastened to the plate using masking tape. A Microplate Genie Digital (Scientific Industries, USA) was used for agitation. This microtitre plate vortex has an orbit of 1 mm with accurate speed control. The rotation speed was confirmed using a Voltcraft DT-10L digital tachometer (Conrad, Germany). The entire setup was placed in a humidity-controlled incubator set to 70% RH and 30°C. The evaporation rate was around 8 μL per well per day.

### Image analysis

Image analysis was performed as described by whole slide imaging combined with automated image analysis, using the SParticle algortihm that was developed specifically as Plugin for imageJ (Willemse 2017). In short, 100 μl sample was transferred to a glass microscope slide and covered by a 24×60 cover slip. The slide was mounted in an Axio Observer (Zeiss, Germany) equipped with an automated XY-stage, which allowed whole slide imaging using a 10x objective. The imageJ plugin for automated image analysis optimized for *Streptomyces* liquid morphology was used to obtain both the maximum Feret length and a shape description for circularity (Stojmenovic 2013) of each mycelial fragment or pellet found in the sample. Incorrectly analysed pellets (e.g. out-of-focus mycelia) were removed manually. Further data processing and visualization was done in Microsoft Excel.

### Tyrosinase acitivity measurement

Tyrosinase activity was measured by the conversion over time l-3,4-dihydroxyphenylalanine spectrophotometrically at a wavelength of 475 nm, as described (van Wezel 2006).

### Actinorhodin quantification

The production of actinorhodin by *S. coelicolor* was determined as follows. Culture supernatant (40 μl) was treated with 0.5 μl 5 M HCl to pH 2 to 3, extracted with a 0.5 volume of methanol-chloroform (1:1), and centrifuged at 5,000 rpm for 10 min. The concentration was calculated from the A_542_ (ε542, 18,600).

## RESULTS

### The morphology of *S. lividans* in shake flasks

To scale down the culture volume, while retaining the morphology, we aimed at replicating key morphological parameters of the liquid-based growth of *S. lividans* in a shake flask, such as pellet formation and fragmentation. We applied the SParticle plugin for ImageJ to quantify the size and shape distribution of the pellets via whole-slide image analysis (Willemse 2017).

As a reference, the morphological characteristics of shake flask cultures were investigated. Around 500 aggregates were analysed from three separate 24 h shake flask-grown cultures, corresponding to the end of the exponential growth phase, which roughly corresponds to the moment of antibiotic production initiation (Nieselt 2010). Previous work comparing the maximum length of pellets revealed two different mycelial populations of *S. lividans*, one forming larger and one smaller pellets (van Veluw 2012). This separation is even more apparent when the shape of a particle, measured as the circularity, is taken into account. This revealed two distinct clusters of particles that not only differ in size, but also in shape (Fig. 1, scatter plot). One population had pellets with similar lengths of around 200 μm, but with a wide standard deviation in circularity (Fig. 1D, falling within the orange dotter oval and Fig. 1C). The other, representing the majority of pellets, were found in clusters and were more regularly shaped, often slightly oval and with homogenous density (Fig. 1C and Fig. 1D, purple striped oval). Pellets with a large size frequently lost their structural integrity,presumably as they were in the process of disintegration (Fig. 1B). Because of the effects of pellets on production and regulation it is important to capture all of these morphological characteristics when scaling down.

**Figure 1.**
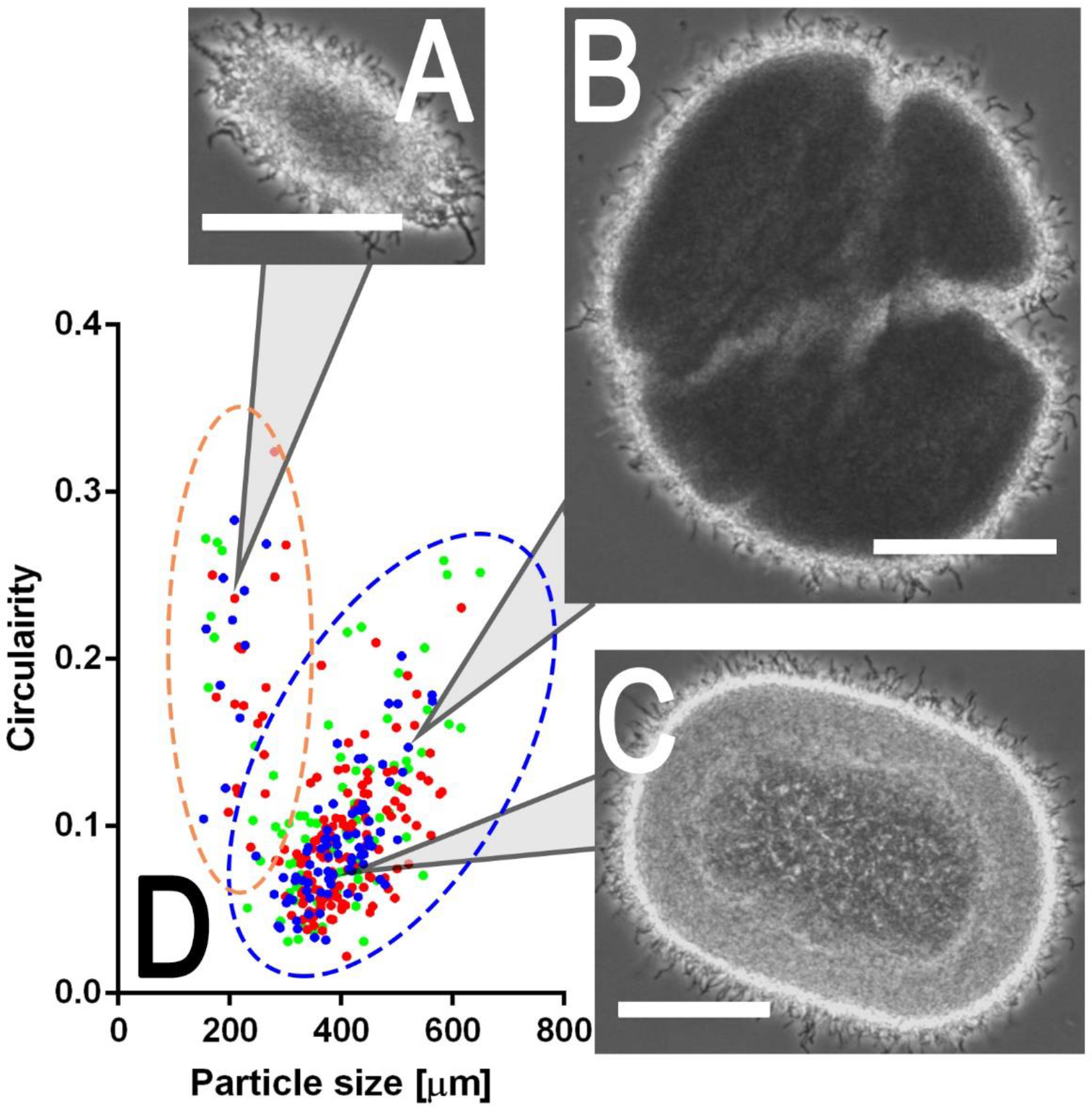
Morphological characterization of *S. lividans* in shake flask cultures. Spores of *S. lividans* 66 were inoculated at 10^6^ CFU/ml into a shake flask (equipped with a coiled spring) containing 30 mL TSBS. The culture was grown for 24 h in an orbital shaker set to 30°C. Three 100 μL samples from two independent cultures were taken at 24 h and subjected to image analysis to obtain the maximum length and circularity of each distinguishable mycelial aggregate. Light micrographs A, B and C represent the three archetypes of the pellet morphologies seen in the culture, and corresponds to the indicated locations in the particle size (x-axes) and circularity (y-axes) scatterplot (D). Two distinct clusters can be distinguished (yellow and blue dotted circles)The data is obtained from three biological replicates. Bar, 100 μm.

### Dependency of agitation rate on the morphology in micro-cultures

The insight that pellet formation is mostly the result of the hydrodynamic forces and/or the supply of sufficient oxygen, often described by the power dissipation (P) and the k_L_a, prompted new experiments to match the environmental properties with those found in shake flasks. Both of these properties can be changed with (and are linked to) the agitation rate. A digital vortex, designed for microtitre plate (MTP) mixing, with its variable speed setting allowed the study of morphology in relation to the agitation speed. This was used to establish whether a population could be obtained with morphological characteristics similar to those found in larger scale cultures.

In 100 μl MTP cultures and at low agitation rates, the mycelia failed to aggregate into the dense pellets normally observed in shake flasks, showing instead a more irregularly shaped open morphology (Fig. 2A and 2G). Apart from the lower density, the average length of the mycelia was around 480 μm, but with a broad distribution; particle sizes ranged from 80 μm to huge aggregates of up to 1500 μm (Table 1). The low number of pellets per well may be caused by extensive spore aggregation and/ or lack of mycelial fragmentation under these growth conditions.

**Figure 2.**
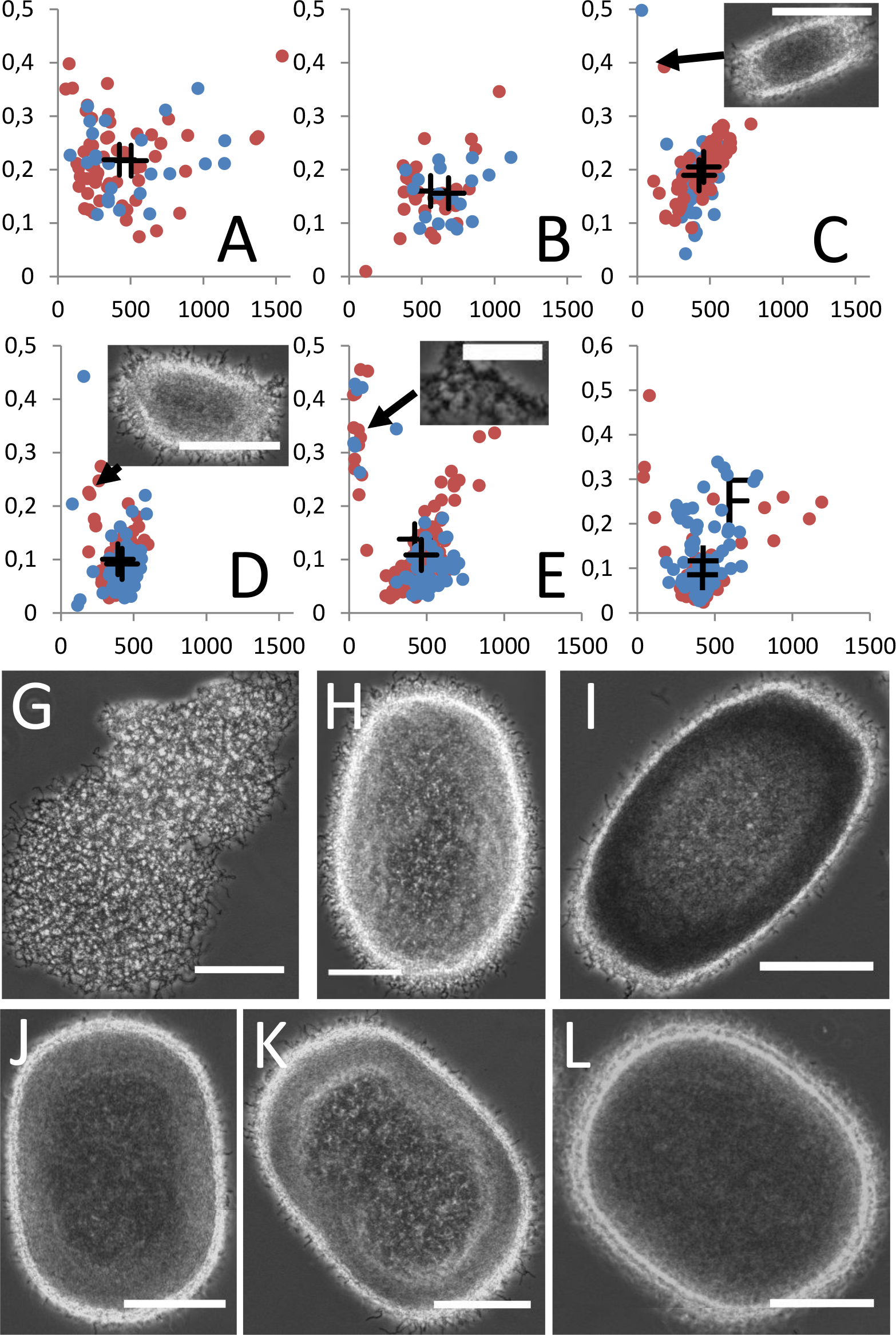
Growth of *S. lividans* in 100 μL MTP cultivation at different agitation rates. Spores of *S. lividans* 66 were inoculated at 10^6^ CFU/ml into 100 μL in a 96 well MTP with V-shaped bottom. The MTPs were agitated using a digital MTP vortex, which was set in a humidified incubator with temperature set to 30°C. The agitation rate was changed between experiments ranging from 800 rpm (A,G), 1000 rpm (B,H), 1200 rpm (C,I), 1400 rpm (D,J), 1600 rpm (E,K) and 1800 rpm (F,L) and the effects on morphology of each aggregate after 24 h of cultivation was analysed in respect to its circularity (y-axes) and maximum length (x-axes) and are displayed in a scatter plot (A,B,C,D,E,F). Agitation rates were analysed twice (replicates presented in blue and red) and the centroids calculated (black crosses). C,D,E: image showing an aggregate of the smaller population. G,H,I,J,K,L: examples of a typical pellet found near the centroid. Scale bars: 50 μm (E) or 100 μm (all other images).

**Table 1.**
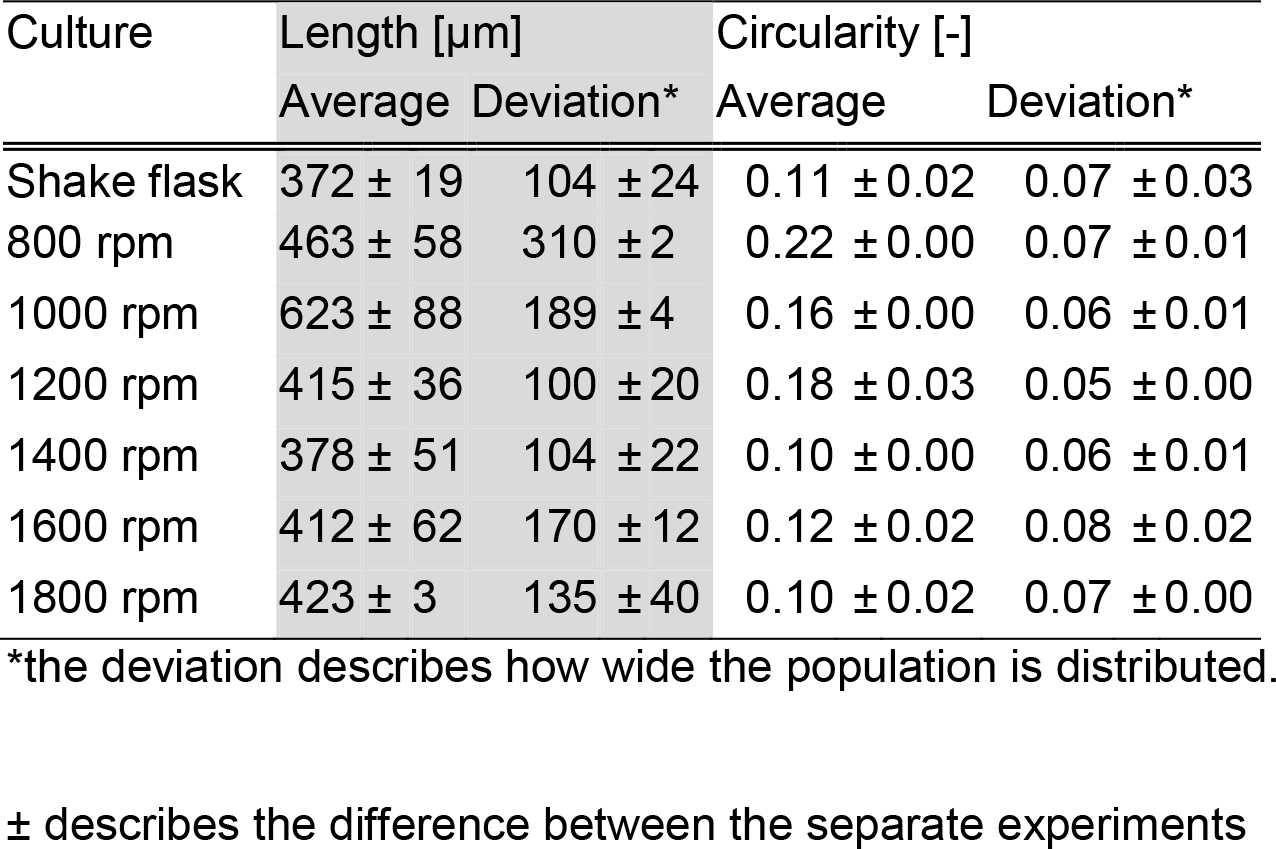
Average length and circularity of the population of mycelial aggregates under different growth conditions.

At 1000 rpm the mycelia formed denser pellets, typical of *S. lividans* grown in shake flasks or in the fermenter (Fig. 2H). However, the average pellet size of around 600 μm and the significantly smaller number of particles per culture volume suggest that the hydrodynamic forces in de micro-cultures were weaker than in shake flask-grown cultures (Fig. 2B). At 1200 rpm the average pellet size approached that found in shake flasks, but the pellets had a more elongated, oval-shaped morphology, with an average roundness close to 0.2 (Fig. 2C and 2I). At this agitation rate a few pellets of the vertical cluster appeared, indicating that the shear stress was sufficiently high to induce fragmentation (Fig. 2C, inset). Increasing the mixing rate further to 1400 rpm lowered the circularity to the desired value of 0.1, and now the average pellet length closely resembled that obtained in shake-flask cultures (Fig. 2D and 2J). Also the distribution of the pellet population closely resembled those formed in shake flasks, including the occurrence of the population of smaller oval-shaped pellets (Fig. 2D, inset). When the agitation was further increased to 1600 rpm the morphological characteristics of the cultures showed again an increase in average length and a decrease in circularity (Fig. 2I and J). Also the Feret diameter of the second population pellets decreased strongly, indicating that the shear stress caused substantial cell damage. Increasing the agitation further to 1800 rpm resulted in a similar trend as seen in 1600 rpm where larger pellets again appeared as part of the population. This may be explained by the culture fluid showing “out of phase” characteristics at high rotations speeds (Büchs 2000; Büchs 2001), which would result in a lower power consumption, and thus lower fragmentation rates and increased average pellet size.

### Production of heterologous enzymes and antibiotics

Mycelia of 24 h old microcultures grown at an agitation speed of 1400 rpm generally had a morphology that was very similar to that observed for mycelia grown in shake-flasks. To analyse how similar the cultures were in terms of their producing capacity, we analysed the production of tyrosinase, which is a good model system for extracellular enzyme production, and was heterologously expressed in *S. lividans* by the introduction of plasmid pIJ703 (van Wezel 2006). A similar amount of active enzyme was produced in shake flasks (200 rpm, 1 inch orbital) and in MTPs (1400RPM, 1 mm orbital), although production started slightly earlier in MTPs (Fig. 3).

**Figure 3.**
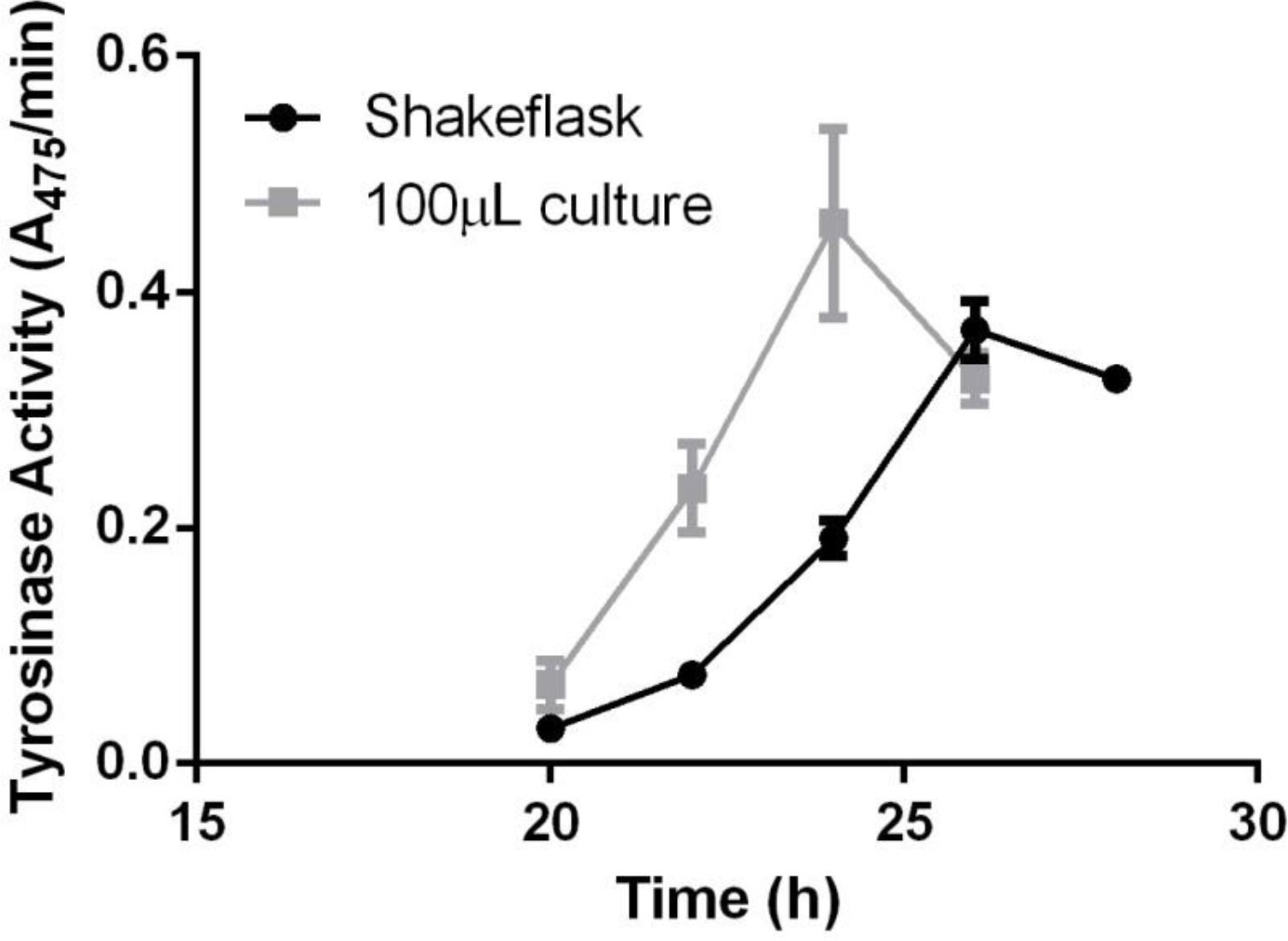
Tyrosinase production by *S. lividans* in shake flasks and 100 μL cultures. Transformants of *S. lividans* 66 heterologously expressing the secreted enzyme tyrosinase from plasmid pIJ703 were grown in TSBS in either shake flasks or V-bottom MTPs. The graph represents the conversion rate of l-3,4-dihydroxyphenylalanine by the culture supernatant, which is indicative of tyrosinase activity. The shake flasks were run in duplicate, while the tyrosinase was measured in three independent wells in the MTP.

To study the effect on antibiotic production, we used *Streptomyces coelicolor* M145 as the model organism, as this strain produces the pigmented polyketide antibiotics actinorhodin and undecylprodigiosin, which are readily assessed spectrophotometrically. For this study, we compared the production of the blue-pigmented actinorhodin between shake flasks and microcultures (Fig. 4). After 48 h of growth both cultures had produced comparable amounts of actinorhodin, indicating that production of this antibiotic is comparable between the two culturing methods.

**Figure 4.**
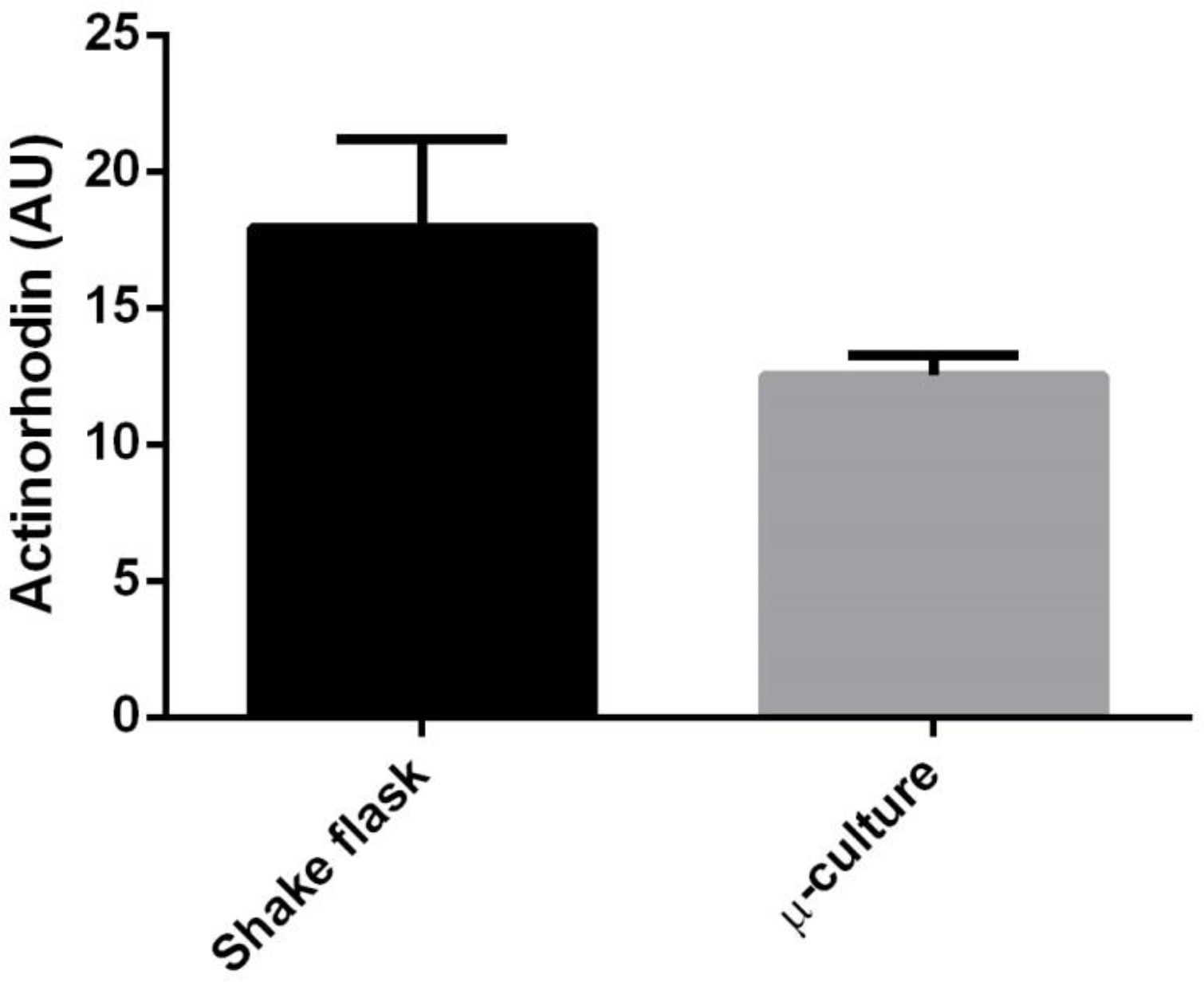
Actinorhodin production by *S. coelicolor* after 48 h of growth. M145 was cultivated in minimal media for 48 h. The shake flasks were run in duplicate, while antibiotic production was measured in three different wells in the MTP. Actinorhodin was extracted by chloroform/methanol and measured spectrophotometrically at 542 nm. The average amount of actinorhodin (in arbitrary units) concentration and the SEM of three independent cultures are shown.

## DISCUSSION

High-throughput screening of actinobacteria for natural products or enzymes typically takes place in micro-scale liquid-grown cultures in an MTP-based setup. The alternative is solid-grown cultures, but it is very difficult to translate growth conditions from solid- to liquid-grown cultures. A drawback of screening of actinobacteria in submerged cultures is the formation of large mycelial networks, which show flocculation or attachment to abiotic surfaces and are associated with slow growth (van Dissel 2014). Additionally, cultures tend to be highly heterogeneous due to the large surface area of the mycelial clumps. Recently, we showed that aggregation of germlings increases culture heterogeneity (Zacchetti 2016). Because heterogeneity creates a distribution of morphologies, all contributing to production differently (van Veluw 2012; Martin and Bushell 1996), a large population is often required to maintain reproducibility. As heterogeneity is influenced by environmental parameters, careful control is needed to mimic the morphology of a shake flask in small-scale cultivation platform.

Growth in small volumes typically favours pellet formation, and genetically engineered strains have been developed that result in dispersed growth, via over-expression of the cell division activator gene *ssgA* (van Wezel 2006; Traag and van Wezel 2008; van Dissel 2015). However, such genetic manipulation often has major consequences for product formation. Recently, the Biolector system, allows parallel growth of 48 cultures in a MTP with around 1 mL volume, was successfully adapted for growth and screening of streptomycetes (Koepff 2017). The authors obtained promising results, obtaining growth parameters that could be compared to thse seen in 1 L cultures. In this study, we show that streptomycetes can be successfully cultivated even in 100 μl microcultures, without the use of specialized equipment, while maintaining the same morphology as in large shake flasks. Our data show that the distribution of a heterogeneous mycelial population is highly dependent on the agitation rate in 96-well MTPs. Especially at insufficient mixing rates the mycelia failed to aggregate into typical pellet structures. Possibly this is in part the result of insufficient oxygen supply. The relationship between oxygen supply and morphology is not well understood, but preliminary experiments where the oxygen supply was limited in a shake flask by reducing the gas exchange, resulted in pellets with a reduced density similar to what was found in poorly agitated MTPs (DvD and GPvW, unpublished results). Although the k_L_a was not measured in this study, initial calculations using equations for orbital mixing (Seletzky 2007) showed that the oxygen transfer could be lower than adequate, with a k_L_a as low as 40 h^-1^, for a mixing rate of 800 rpm. This low value is suggestive of oxygen limitation as the cause of the morphology observed at low agitation rates and that in part the change in morphology by increased agitation is the result of an increased oxygen supply. While these observations are indicative of oxygen limitation as determining factor for mycelial morphology, oxygen transfer and hydrodynamic stress are coupled processes for orbital shaken cultivation methods (and to some extent also for bioreactors). At least for pristinamycin production hydrodynamic stress, described as the power input, was more descriptive for both pellet morphology and production levels (Mehmood 2012). Downscaling to 100 μl is feasible for the filamentous *Streptomyces*, even if it aggregates into dense pellets. How precisely agitation affects morphogenesis in MTP plates is as yet unclear and requires further study.

Matching the environment, including the physical hydrodynamic forces that determine the morphology is due to its complex nature a difficult task. Our study also illustrates the utility of image analysis to quantify the morphology and assist in the down-scaling process. It follows that detailed comparison of mycelial morphologies by image analysis allows selection of the appropriate culturing conditions to obtain a preferred average pellet size and structure. Comparison of maximal pellet length and circularity provides more detailed insights into the exact morphology of the pellets, which aids the down-scaling process. Besides providing the option of medium- to high-throughput screening, the ability to grow *Streptomyces* with a native morphology on a very small scale also allows studies that involve for example the addition of expensive or low abundance chemicals or enzymes.

## CONCLUSION

The complex morphology displayed by filamentous actinobacteria in liquid-grown cultures greatly influences their productivity. Screening these bacteria for new therapeutic agents in an MTP-based setup without affecting normal growth and morphology would be a major advantage. This is particularly important in the light of upscaling, so as to maximise the chance that productivity is maintained. We have been able to translate growth and morphology from shake flasks to 100 μL micro cultures by carefully tuning the rate of agitation. The resulting growth and average pellet size in standard HTS-compatible MTPs was reproducibly comparable to those in larger scale cultures, which is an important contribution to the state of the art.

## Declarations

### Authors’ contributions

D van Dissel and GP van Wezel defined the research theme and designed the experiments. D van Dissel carried out the laboratory experiments. D van Dissel and GP van Wezel analysed the data, interpreted the results and prepared this manuscript. Both authors approved the final manuscript.

## Acknowledgements

The work was supported by VICI grant 10379 from the Netherlands Organization for Scientific Research (NWO).

## List of abbreviations

CFU: colony forming units
HTS: high throughput screening
MTP: microtitre plate
rpm: rotations per minute
k_L_a: specific oxygen transfer coefficient
P: power dissipation

## References

Anné J, Maldonado B, Van Impe J, Van Mellaert L, Bernaerts K (2012) Recombinant protein production and streptomycetes. J Biotechnol 158: 159–167.

Barka EA, Vatsa P, Sanchez L, Gavaut-Vaillant N, Jacquard C, Meier-Kolthoff J, Klenk HP, Clément C, Oudouch Y, van Wezel GP (2016) Taxonomy, physiology, and natural products of the *Actinobacteria*. Microbiol Mol Biol Rev 80: 1–43.

Bérdy J (2005) Bioactive microbial metabolites. J Antibiot (Tokyo) 58: 1–26.

Büchs J, Lotter S, Milbradt C (2001) Out-of-phase operating conditions, a hitherto unknown phenomenon in shaking bioreactors. Biochem Eng J 7: 135–141.

Büchs J, Maier U, Milbradt C, Zoels B (2000) Power consumption in shaking flasks on rotary shaking machines: II. Nondimensional description of specific power consumption and flow regimes in unbaffled flasks at elevated liquid viscosity. Biotechnol Bioeng 68: 594–601.

Celler K, Picioreanu C, van Loosdrecht MC, van Wezel GP (2012) Structured morphological modeling as a framework for rational strain design of Streptomyces species. Antonie Van Leeuwenhoek 102: 409–423.

Chater KF, Losick R (1997) Mycelial life style of *Streptomyces coelicolor* A3(2) and its relatives. In: Shapiro JA, Dworkin M (eds) Bacteria as multicellular organisms. Oxford University Press, New York, pp 149–182.

Claessen D, Rozen DE, Kuipers OP, Sogaard-Andersen L, van Wezel GP (2014) Bacterial solutions to multicellularity: a tale of biofilms, filaments and fruiting bodies. Nat Rev Microbiol 12: 115–124.

Cruz-Morales P,Vijgenboom E, Iruegas-Bocardo F, Girard G, Yanez-Guerra LA, Ramos-Aboites HE, Pernodet JL, Anne J, van Wezel GP, Barona-Gomez F (2013) The genome sequence of *Streptomyces lividans* 66 reveals a novel tRNA-dependent peptide biosynthetic system within a metal-related genomic island. Genome Biol Evol 5: 1165–1175.

Dobson L, O’Cleirigh C, O’Shea D (2008) The influence of morphology on geldanamycin production in submerged fermentations of *Streptomyces hygroscopicus* var. *geldanus*. Appl Microbiol Biotechnol 79: 859–866.

Duetz WA, Ruedi L, Hermann R, O’Connor K, Buchs J, Witholt B (2000) Methods for intense aeration, growth, storage, and replication of bacterial strains in microtiter plates. Appl Environ Microbiol 66: 2641–2646.

Hopwood DA (2007) Streptomyces in nature and medicine: the antibiotic makers. Oxford University Press, New York.

Huber R, Ritter D, Hering T, Hillmer AK, Kensy F, Muller C, Wang L, Buchs J (2009) Robo-Lector - a novel platform for automated high-throughput cultivations in microtiter plates with high information content. Microb Cell Fact 8: 42.

Jakimowicz D, van Wezel GP (2012) Cell division and DNA segregation in *Streptomyces:* how to build a septum in the middle of nowhere? Mol Microbiol 85: 393–404.

Katz E, Thompson CJ, Hopwood DA (1983) Cloning and Expression of the Tyrosinase Gene from *Streptomyces antibioticus* in *Streptomyces lividans*. J Gen Microbiol 129: 2703–2714.

Kieser T, Bibb MJ, Buttner MJ, Chater KF, Hopwood DA (2000) Practical streptomyces genetics.

Koepff J, Keller M, Tsolis KC, Busche T, Ruckert C, Hamed MB, Anne J, Kalinowski J, Wiechert W, Economou A, Oldiges M (2017) Fast and reliable strain characterization of *Streptomyces lividans* through micro-scale cultivation. Biotechnol Bioeng.

Kolter R, van Wezel GP (2016) Goodbye to brute force in antibiotic discovery? Nat Microbiol 1: 15020.

Liman R, Facey PD, van Keulen G, Dyson PJ, Del Sol R (2013) A laterally acquired galactose oxidase-like gene is required for aerial development during osmotic stress in *Streptomyces coelicolor*. PLoS ONE 8: e54112.

Long Q, Liu X, Yang Y, Li L, Harvey L, McNeil B, Bai Z (2014) The development and application of high throughput cultivation technology in bioprocess development. J Biotechnol 192: 323–338.

Martin SM, Bushell ME (1996) Effect of hyphal micromorphology on bioreactor performance of antibiotic-producing *Saccharopolyspora erythraea* cultures. Microbiol 142: 1783–1788.

McCormick JR (2009) Cell division is dispensable but not irrelevant in *Streptomyces*. Curr Opin Biotechnol 12: 689–698.

Mehmood N, Olmos E, Goergen J-L, Blanchard F, Marchal P, Klöckner W, Büchs J, Delaunay S (2012) Decoupling of oxygen transfer and power dissipation for the study of the production of pristinamycins by *Streptomyces pristinaespiralis* in shaking flasks. Biochem Eng J 68: 25–33.

Minas W, Bailey JE, Duetz W (2000) Streptomycetes in micro-cultures: Growth, production of secondary metabolites, and storage and retrieval in the 96–well format. Antonie van Leeuwenhoek 78: 297–305.

Nieselt K, Battke F, Herbig A, Bruheim P, Wentzel A, Jakobsen ØM, Sletta H, Alam MT, Merlo ME, Moore J (2010) The dynamic architecture of the metabolic switch in *Streptomyces coelicolor*. BMC genomics 11: 10.

Ohta N, Park YS, Yahiro K, Okabe M (1995) Comparison of neomycin production from *Streptomyces fradiae* cultivation using soybean oil as the sole carbon source in an air-lift bioreactor and a stirred-tank reactor. J Ferm and Bioeng 79: 443–448.

Olmos E, Mehmood N, Husein LH, Goergen J-L, Fick M, Delaunay S (2013) Effects of bioreactor hydrodynamics on the physiology of *Streptomyces*. Bioproc Biosyst Eng 36: 259–272.

Petrus ML, Vijgenboom E, Chaplin AK, Worrall JA, van Wezel GP, Claessen D (2016) The DyP-type peroxidase DtpA is a Tat-substrate required for GlxA maturation and morphogenesis in *Streptomyces*. Open Biol 6.

Rohe P, Venkanna D, Kleine B, Freudl R, Oldiges M (2012) An automated workflow for enhancing microbial bioprocess optimization on a novel microbioreactor platform. Microb Cell Fact 11: 144.

Roubos JA, Krabben P, Luiten RGM, Verbruggen HB, Heijnen JJ (2001) A quantitative approach to characterizing cell lysis caused by mechanical agitation of *Streptomyces clavuligerus*. Biotechnol Progr 17: 336–347.

Seletzky JM, Noak U, Fricke J, Welk E, Eberhard W, Knocke C, Büchs J (2007) Scale-up from shake flasks to fermenters in batch and continuous mode with *Corynebacterium glutamicum* on lactic acid based on oxygen transfer and pH. Biotechnol Bioeng: 800–811.

Siebenberg S, Bapat PM, Lantz AE, Gust B, Heide L (2010) Reducing the variability of antibiotic production in *Streptomyces* by cultivation in 24-square deepwell plates. J Biosci Bioeng 109: 230–234.

Sohoni SV, Bapat PM, Lantz AE (2012) Robust, small-scale cultivation platform for *Streptomyces coelicolor*. Microb Cell Fact 11: 9.

Stojmenovic M, Jevremovic A, Nayak A Fast iris detection via shape based circularity. In: Industrial Electronics and Applications (ICIEA), 2013 8th IEEE Conference on, 2013. IEEE, pp 747–752.

Tamura S, Park Y, Toriyama M, Okabe M (1997) Change of mycelial morphology in tylosin production by batch culture of *Streptomyces fradiae* under various shear conditions. J Ferment Bioeng 83: 523–528.

Traag BA, van Wezel GP (2008) The SsgA-like proteins in actinomycetes: small proteins up to a big task. Antonie Van Leeuwenhoek 94: 85–97.

van Dissel D, Claessen D, Roth M, van Wezel GP (2015) A novel locus for mycelial aggregation forms a gateway to improved *Streptomyces* cell factories. Microb Cell Fact: In press.

van Dissel D, Claessen D, van Wezel GP (2014) Morphogenesis of *Streptomyces* in submerged cultures. Adv Appl Microbiol 89: 1–45.

van Veluw GJ, Petrus ML, Gubbens J, de Graaf R, de Jong IP, van Wezel GP, Wosten HA, Claessen D (2012) Analysis of two distinct mycelial populations in liquid-grown *Streptomyces* cultures using a flow cytometry-based proteomics approach. Appl Microbiol Biotechnol 96: 1301–1312.

van Wezel GP, Krabben P, Traag BA, Keijser BJ, Kerste R, Vijgenboom E, Heijnen JJ, Kraal B (2006) Unlocking *Streptomyces* spp. for use as sustainable industrial production platforms by morphological engineering. Appl Environ Microbiol 72: 5283–5288.

van Wezel GP, McKenzie NL, Nodwell JR (2009) Chapter 5. Applying the genetics of secondary metabolism in model actinomycetes to the discovery of new antibiotics. Methods Enzymol 458: 117–141.

Vrancken K, Anne J (2009) Secretory production of recombinant proteins by Streptomyces. Future Microbiol 4: 181–188.

Willemse J, Büke F, van Dissel D, Grevink S, Claessen D, van Wezel GP (2017) SParticle, an algorithm for the analysis of filamentous microorganisms in submerged cultures. BioRXiv: 159475.

Wucherpfennig T, Kiep KA, Driouch H, Wittmann C, Krull R (2010) Morphology and rheology in filamentous cultivations. In, vol Volume 72. Academic Press, pp 89–136.

Zacchetti B, Willemse J, Recter B, van Dissel D, van Wezel GP, Wosten HA, Claessen D (2016) Aggregation of germlings is a major contributing factor towards mycelial heterogeneity of *Streptomyces*. Sci Rep 6: 27045.

